# A lethal mouse model of Oz virus infection reveals hepatic involvement and enables evaluation of antiviral and vaccine efficacy

**DOI:** 10.64898/2026.01.29.702403

**Authors:** Rio Harada, Cassia Sousa Moraes, Atsuko Inoue, Chilekwa Frances Kabamba, Koshiro Tabata, Motohito Goto, Riichi Takahashi, Joshua W. Kranrod, Shuichi Osawa, Aiko Ohnuma, Saito Shinji, Gabriel Gonzalez, Noriko Nagata, Mamoru Ito, Makoto Suematsu, Hideki Hasegawa, William W. Hall, Hirofumi Sawa, Yukari Itakura

## Abstract

Oz virus (OZV), a member of the genus *Thogotovirus* in the family *Orthomyxoviridae*, is an emerging tick-borne virus reported in Japan. A fatal human case and seroepidemiological evidence of widespread exposure among wild animals and humans suggest its potential public health significance. However, no animal models suitable for pathogenic studies or evaluation of countermeasures are available for OZV. Here, we have established a lethal mouse model of OZV infection using cell-adapted virus and mice lacking type I interferon signaling (B6 Ifnar1 KO mice). OZV infection resulted in 100% mortality and was characterized by robust viral replication in the liver and spleen, severe hepatitis, and acute liver injury. Using this model, we also demonstrated that oral administration of T-705, an antiviral drug widely used against RNA viruses, as well as immunization with an inactivated whole virus particle vaccine, protected B6 Ifnar1 KO mice from lethal OZV infection by mitigating the acute hepatitis. The present study provides critical insights into OZV pathogenesis and establishes a practical in vivo platform for the development of countermeasures against OZV infection.

**Significance statement:** Emerging tick-borne viruses pose a growing public health concern, yet progress in understanding their pathogenesis and in developing countermeasures is often limited by the lack of relevant animal models. Oz virus (OZV), a recently identified thogotovirus associated with a fatal human case, exemplifies this challenge. Here, we establish a lethal mouse model of OZV infection that reveals pronounced hepatic involvement as a central pathological feature. Using this model, we demonstrate effective protection by both an antiviral drug and an inactivated vaccine against lethal OZV challenge. This study provides a practical in vivo platform for investigating OZV pathogenesis and for accelerating the development of medical countermeasures against this emerging tick-borne virus.

## Introduction

Oz virus (OZV) is a six-segmented, negative-sense, single-stranded RNA virus belonging to the genus *Thogotovirus* in the family *Orthomyxoviridae* [1]. Thogotoviruses are tick-borne viruses with a global distribution [2–4] and infect a broad range of vertebrate hosts [2,5– 8]. Serological evidence of infection has been documented in domestic and wild animals, including cattle, sheep, raccoons, rabbits, and deer, as well as humans [2,5–8]. Several members of this genus, including Thogoto virus (THOV), Dhori virus (DHOV), and Bourbon virus (BRBV), are known to cause febrile illness, encephalitis, and even fatal outcomes in humans [2,9,10] underscoring their public health relevance despite the limited number of reported cases.

OZV was first isolated from hard ticks, *Amblyomma testudinarium*, in Ehime, Japan, in 2018 [11]. To date, one fatal human case of OZV infection, associated with myocarditis, was reported in Ibaraki, Japan, in 2022 [12], demonstrating its pathogenic potential in humans. Subsequent seroepidemiological studies have revealed the wide distribution of OZV in wild animals, including Japanese macaques, wild boars, black bears, and deer, as well as humans [13– 16]. Moreover, repeated detection of OZV in ticks and viral genome fragments in wild animals across multiple regions of Japan [15–17] indicates sustained circulation and ongoing zoonotic transmission risks.

Despite these concerns, virological and pathogenic characterization of OZV, as well as the development of effective countermeasures, have been hindered by the lack of a suitable animal models. Although OZV can cause lethality following intracerebral inoculation in suckling mice [11], peripheral infection of adult immunocompetent mice does not result in detectable virus replication or disease [18]. These observations suggest that OZV infection is efficiently regulated by innate immune defenses. Consistent with this notion, BRBV, a thogotovirus closely related to OZV, exhibits strict dependence on impaired interferon (IFN) signaling to establish productive infection in vivo, and disruption of type I IFN pathways is essential for the development of lethal mouse models suitable for elucidation of pathogenesis and evaluation of countermeasures [19,20].

Hypothesizing that OZV pathogenicity is likewise constrained by type I IFN-mediated antiviral responses, here, we aimed to establish a lethal mouse model of OZV infection using mice lacking type I IFN receptors, which have been used as animal models in various virus studies [10,19,20]. Employing this model of OZV infection, we demonstrate that OZV infection results in reproducible lethality, with robust viral replication in the liver and spleen, causing severe hepatic injury. Furthermore, we validate the utility of this model by demonstrating complete protection against lethal challenges by both the antiviral agent T-705 and an inactivated whole-virus vaccine. Together, this lethal mouse model provides a valuable in vivo platform for investigating OZV pathogenesis and for developing countermeasures against this emerging tick-borne virus infection.

## Results

### Generation and characterization of cell-adapted OZV

To establish a mouse infection model for OZV, we first generated a cell-adapted virus that efficiently replicates in vitro. After the serial passage of the OZV Ibaraki/O10-S/2022 strain, the clinical isolate from the human fatal case [12], in Vero E6 cells, the replication efficacy of the virus significantly increased with an approximate 600-fold increase in the highest yield between passage 6 and passage 17 (OZV-P17) (S1 Fig).

### Lack of productive infection of cell-adapted OZV in BALB/c mice

To assess the pathogenic potential of OZV-P17 in immunocompetent mice, 3-week-old, 8-week-old, and aged BALB/c mice were inoculated with 10^6^ plaque-forming units (PFU) of OZV-P17 via subcutaneous (s.c.), intraperitoneal (i.p.), or intradermal routes and monitored for 20 days post-infection (dpi). None of the mice exhibited any symptoms or body weight loss during the observation period (S2A Fig). Nevertheless, neutralizing antibodies were detected in all animals (S2B Fig), indicating that asymptomatic infection induced an immune response. Viral replication at 1, 2, 4, 7, 10, and 13 dpi was assessed in the brain, heart, lung, liver, spleen, and serum of OZV-P17-infected 3-week-old BALB/c mice. Infectious virus was not detected in any organs at any time points (data not shown), suggesting that OZV-P17 did not establish a productive infection in immunocompetent mice.

### Establishment of a lethal OZV infection model in B6 Ifnar1 KO mice

Given the resistance of immunocompetent mice to OZV infection, we next generated B6 Ifnar1 KO mice (S3 Fig), which lack type I IFN receptors, and evaluated the susceptibility of the mice. Mice were inoculated with 10^2^, 10^3^, 10^4^, 10^5^, or 10^6^ PFU of OZV-P17 via i.p. or s.c. route, and body weight and survival were monitored for 16 days after infection. In the i.p.-infected groups, most mice exhibited body weight loss beginning at 2 dpi and succumbed to infection between 7 and 8 dpi at several inoculation doses (Fig 1A and S4A Fig). The s.c.-infected groups showed a delayed onset of body weight loss starting at 4 dpi, with death occurring between 7 and 10 dpi (Fig 1B and S4B Fig). The mortality rate ranged from 67% to 100% in both inoculation groups (Fig 1). Interestingly, in some higher titer groups (10^5^ PFU i.p. and 10^5^–10^6^ PFU s.c.), one or two mice recovered from transient body weight loss and survived. Based on consistency and reproducibility of lethality, i.p. inoculation of B6 Ifnar1 KO mice with 10^4^ PFU of OZV-P17 was selected as the standardized lethal OZV infection mouse model in subsequent experiments.

**Fig 1.**
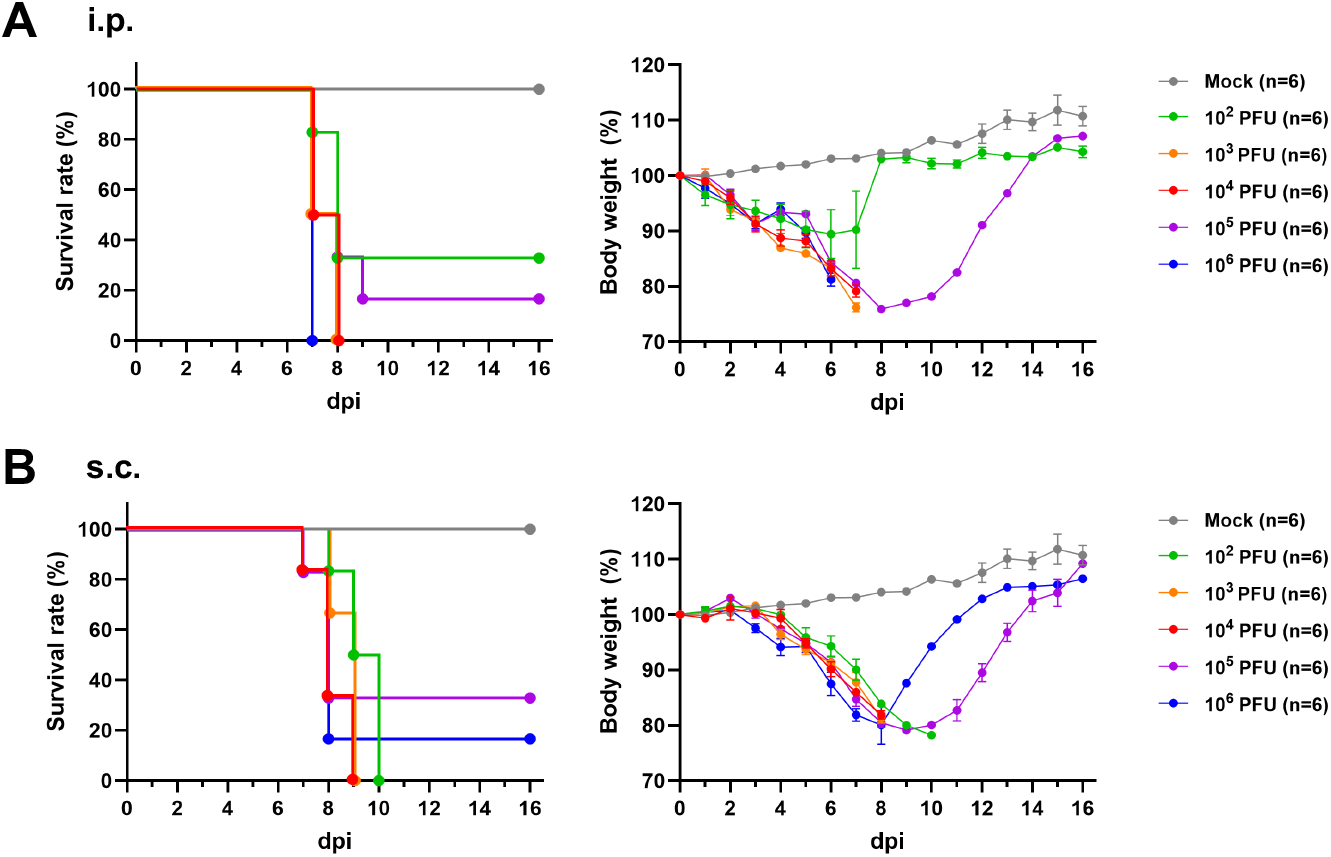
Survival and body weight changes of B6 Ifnar1 KO mice infected with OZV-P17. B6 Ifnar1 KO mice were inoculated with 10^2^, 10^3^, 10^4^, 10^5^, or 10^6^ PFU of OZV-P17 (A) intraperitoneally, or (B) subcutaneously. The same dataset is shown for the mock group in (A) and (B). Survival and body weight changes were monitored daily for 16 days. Data are presented as means ± standard deviation (SD).

### Hepatotropic infection and inflammatory pathology of OZV in B6 Ifnar1 KO mice

To investigate the pathogenesis of OZV in the lethal mouse model, B6 Ifnar1 KO mice were infected intraperitoneally with 10^4^ PFU of OZV-P17, and organs and sera were collected at 1, 3, and 5 dpi. Robust viral replication was observed in the liver and spleen in a time-dependent manner, with viral RNA reaching 10^8^ copies/ml and infectious viral titer reaching approximately 10^6^ PFU/ml at 5 dpi (Fig 2A and 2B). Moderate viral replication was detected in the lung and kidney, with viral RNA levels of 10^7^ copies/ml and viral titer of 10^4^ PFU/ml at 5 dpi (Fig 2A and 2B). In contrast, viral replication in the brain, heart, and serum remained minimal or undetectable (Fig 2A and 2B). Histopathological analysis revealed focal hepatocellular necrosis in the liver and disruption of follicular structures in the spleen beginning at 3 dpi, with lesion severity increasing by 5 dpi (Fig 2C and 2D; S5 and S6 Figs). The significant increase in the histopathological lesion in the liver was also demonstrated quantitatively (Fig 2E and 2F; S7 Fig). Immunohistochemical analysis revealed localization of viral antigens in the necrotic lesion in the liver, whereas widespread viral antigen distribution was observed throughout the spleen (Fig 2C and 2D; S5 and S6 Figs). No virus-related pathological changes or viral antigen were detected in other examined organs (S8 Fig).

**Fig 2.**
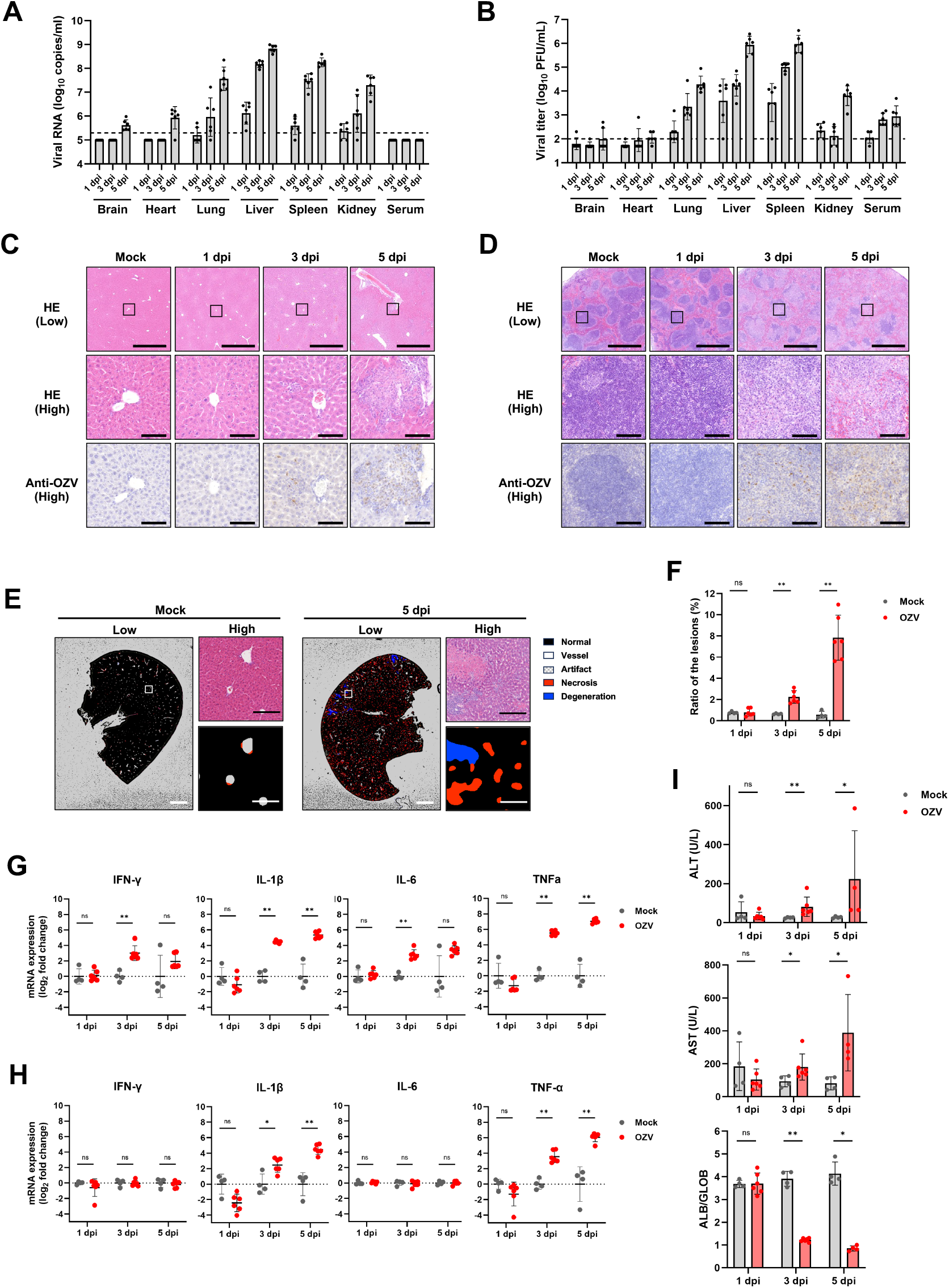
Viral replication and host responses in B6 Ifnar1 KO mice infected with OZV-P17. B6 Ifnar1 KO mice were inoculated intraperitoneally with 10^4^ PFU of OZV-P17. Tissues and sera were collected at 1, 3, and 5 days post infection (dpi). (A, B) Viral replication in each organ was quantified at the indicated time points by (A) RT-qPCR for viral RNA and (B) plaque assay for infectious viral titers. Dotted lines indicate the lower limit of detection. (C, D) Representative images of hematoxylin and eosin (HE) staining and immunohistochemical detection of OZV antigen in the (C) liver and (D) spleen. Scale bars represent 1 mm and 100 µm in the low- and high-magnification images, respectively. (E) Representative images of the histopathological lesion quantification in the liver. Scale bars represent 2 mm and 200 µm in the low- and high-magnification images, respectively. The boxed area in the low-magnification image is shown at high-magnification in panels C–E. (F) Relative area of the histopathological lesions in the liver was quantified. (G, H) Relative mRNA expression levels of inflammatory genes in the (G) liver and (H) spleen at the indicated time points. Gene expression was calculated using the ΔΔCt method and is shown as fold change relative to the mock group. (I) Serum levels of alanine aminotransferase (ALT), aspartate aminotransferase (AST), and albumin-to-globulin ratio (ALB/GLOB). Data are presented as means ± SD, and each dot represents an individual mouse in panels A, B, and F–I. Statistical analyses in panels F–I were performed using the Mann-Whitney U test: *, *P* < 0.05; **, *P* < 0.01.

To characterize host inflammatory responses during lethal OZV infection, mRNA expression levels of IFN-γ, IL-1β, IL-6, and TNF-α were quantified in the liver and spleen using RT-qPCR. These cytokines have been reported to be upregulated in mice lacking type I IFN signaling during various viral infections [21–23]. Consistent with the viral replication and histopathological analysis, expressions of all inflammatory genes were upregulated in the liver (Fig 2G). In contrast, IFN-γ and IL-6 expression were not significantly upregulated in the spleen (Fig 2H). In addition, serum biochemical analysis showed markedly elevated levels of alanine aminotransferase (ALT) and aspartate aminotransferase (AST), liver injury biomarkers (Fig 2I). In contrast, total bilirubin, a biliary tract injury biomarker, remained unchanged (S9 Fig). The albumin-to-globulin ratio decreased, while the levels of total protein were stable, indicating the presence of inflammation and liver dysfunction (Fig 2I and S9 Fig). Decreased levels of alkaline phosphatase and glucose suggested malnutrition may be associated with viral infection (S9 Fig). These results suggested that acute and severe inflammatory responses occurred in the liver. Although a human fatal case presented with severe myocarditis with detection of viral RNA in myocardial cells [12], no efficient viral replication, histopathological lesions, or viral antigen were detected in the hearts of the B6 Ifnar1 KO mice (Fig 2A and 2B; S8 Fig).

### Antiviral efficacy of T-705 in the lethal OZV mouse model

To determine whether the lethal OZV mouse model could be employed in antiviral studies, we assessed the efficacy of T-705, a viral RNA-dependent RNA polymerase inhibitor with broad-spectrum antiviral activity against RNA viruses [24], including BRBV, which is closely related to OZV. OZV-P17-infected B6 Ifnar1 KO mice were orally administered with T-705 at doses of 100 mg/kg/day or 300 mg/kg/day, twice daily from immediately after the infection until 8 dpi (Fig 3A). Body weight and survival were observed for 20 days. All vehicle-treated mice succumbed to infection between 7 and 9 dpi. In contrast, all mice treated with either dose of T-705 survived (Fig 3B) without body weight loss except for the one in the 100 mg/kg/day group showing transient body weight loss and subsequent recovery (S10 Fig). Notably, treatment with 300 mg/kg/day of T-705 completely suppressed viral replication with no detectable infectious virus or viral RNA in all examined organs at 5 dpi (Fig 3C and 3D). Histopathological changes were markedly attenuated in the liver and spleen of the T-705-treated mice (Fig 3E and 3F; S11 Fig). T-705 treatment significantly decreased histopathological lesion formation in the liver compared with vehicle treatment, and localized antigen was undetectable in the treated livers (Fig 3G and 3H; S12 Fig). Although viral antigen was detected in the spleen of the T-705-treated group by immunohistochemical analysis, the number of antigen-positive cells was substantially reduced compared with the vehicle-treated group (Fig 3F and S11 Fig). These results demonstrate the efficacy of T-705 against OZV and the utility of this model for in vivo antiviral evaluation.

**Fig 3.**
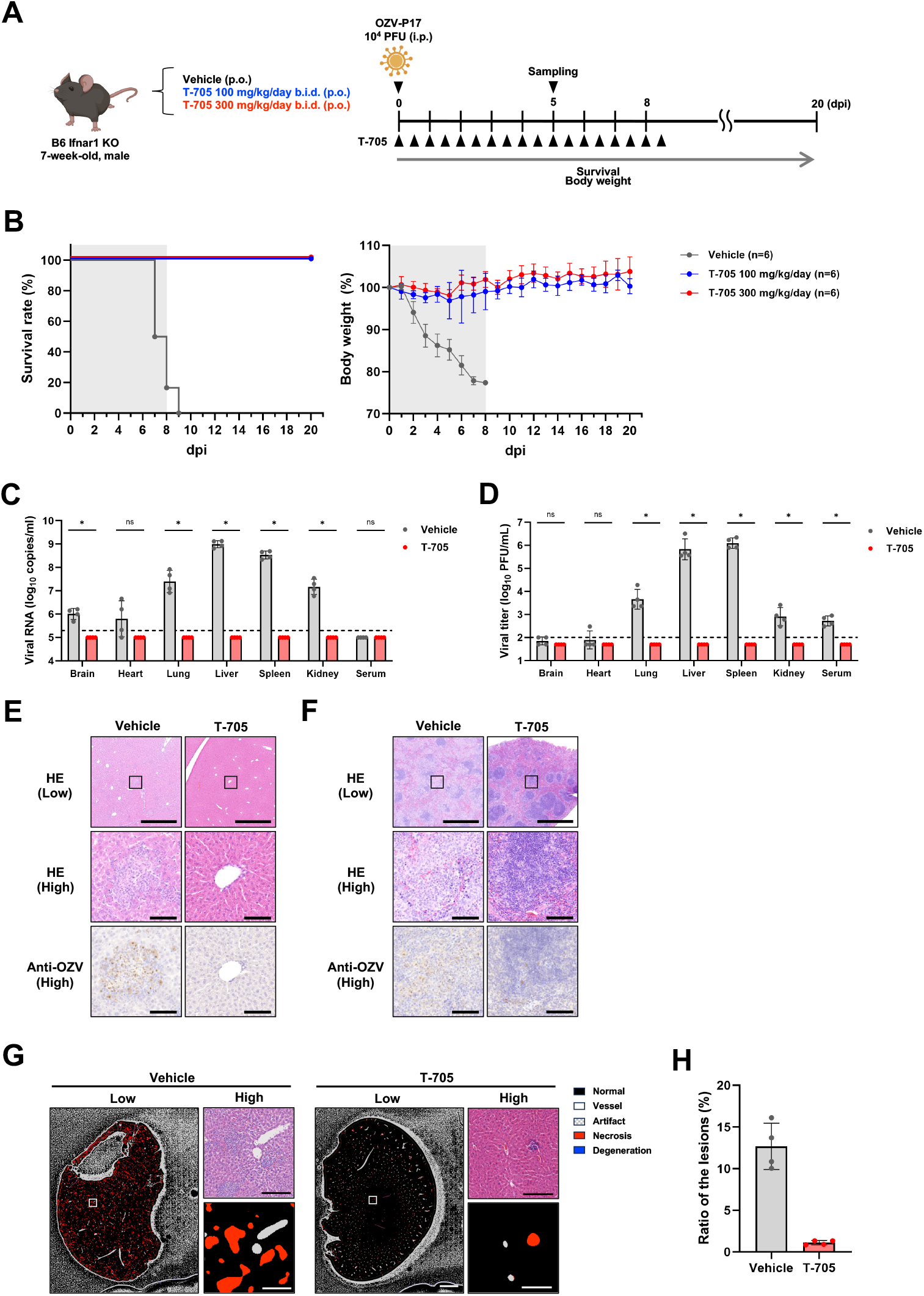
Therapeutic efficacy of T-705 treatment against OZV-P17 infection in B6 Ifnar1 KO mice. (A) Schematic overview of the T-705 treatment experiment. B6 Ifnar1 KO mice were inoculated intraperitoneally with 10^4^ PFU of OZV-P17 and immediately administered orally with 100 or 300 mg/kg/day of T-705, or vehicle for 8 days. (B) Survival and body weight changes were monitored for 20 days. The treatment period is indicated by the gray shaded area. (C, D) Viral replication in liver and spleen was quantified at 5 dpi by (C) RT-qPCR for viral RNA and (D) plaque assay for infectious viral titer. Dotted lines indicate the lower limit of detection. Data are presented as means ± SD. Each dot represents an individual mouse. (E, F) Representative images of HE staining and immunohistochemical detection of OZV in the (E) liver and (F) spleen at 5 dpi. Scale bars represent 1 mm and 100 µm in the low- and high-magnification images, respectively. (G) Representative liver images of the histopathological lesion quantification. Scale bars represent 2 mm and 200 µm in the low- and high-magnification images, respectively. The boxed area in the low-magnification image is shown at high-magnification in panels E–G. (H) The relative area of the histopathological lesions in the liver was quantified. Data are presented as means ± SD in panels B–D, and H, and each dot represents an individual mouse in panels C, D, and H. Statistical analysis in panels C, D, and H was performed using the Mann-Whitney U test: *, *P* < 0.05; **, *P* < 0.01.

### Protective efficacy of an inactivated OZV vaccine in the lethal mouse model

Lastly, we assessed whether the present mouse model could be employed to evaluate the vaccine efficacy against OZV infection with an inactivated whole-virus vaccine (Fig 4A). B6 Ifnar1 KO mice were immunized twice at a 24-day interval with vaccines containing 1.5 µg or 7.5 µg of total viral protein. Both immunization regimens induced robust OZV-specific binding and neutralizing antibody responses. Binding antibody titers were higher in the 7.5 µg group after prime and booster vaccination, whereas neutralizing antibody titers did not differ significantly between the two dose groups (Fig 4B and 4C). At 23 days after booster vaccination, mice were challenged with 10^4^ PFU of OZV-P17. All mice that had received PBS succumbed to infection between 7 and 8 dpi. In contrast, all vaccinated mice survived the lethal challenge (Fig 4D). Transient body weight loss to approximate 80% of baseline was observed in two mice in the 1.5 µg group and one mouse in the 7.5 µg group; however, they recovered and survived (S13 Fig). These results demonstrate that vaccination with inactivated OZV confers robust protective immunity against lethal OZV challenge in this mouse model.

**Fig 4.**
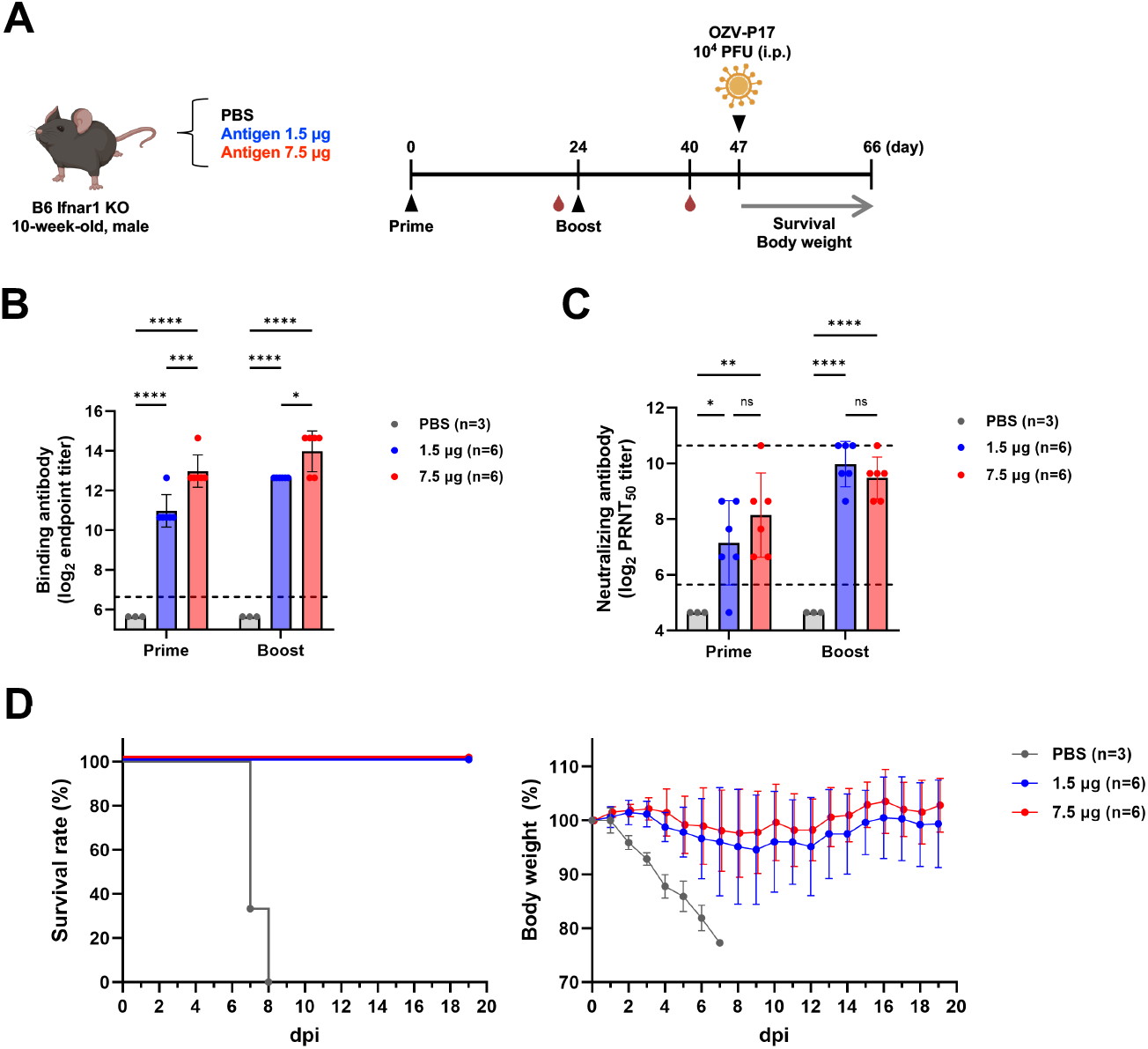
Protective efficacy of vaccination against OZV-P17 infection in B6 Ifnar1 KO mice. (A) Schematic overview of the vaccination and challenge experiments. B6 Ifnar1 KO mice were immunized with 1.5 µg or 7.5 µg of antigen with adjuvant. Mice were challenged intraperitoneally with 10^4^ PFU of OZV-P17. (B, C) OZV-specific (B) binding antibody titers and (C) neutralizing antibody titers in sera were measured after prime and boost immunizations. Dotted lines indicate the lower and upper limit of detection. (D) Survival and body weight changes were monitored for 19 days post-challenge. Data are presented as means ± SD in panels B–D, and each dot represents an individual mouse in B and C. Statistical analysis in panels B and C were performed using two-way ANOVA with Tukey’s multiple comparison test: *, *P* < 0.05; **, *P* < 0.01; ***, *P* < 0.001; ****, *P* < 0.0001.

## Discussion

Previous studies have demonstrated that OZV causes lethal infection in suckling mice but fails to replicate or induce disease in immunocompetent adult mice [11]. Consistent with these findings, we observed no detectable viral replication or pathogenicity in BALB/c mice regardless of age or inoculation route (S2A Fig). Among thogotoviruses, several isolates of THOV and DHOV have been reported to cause lethal infection in immunocompetent mice, whereas Jos virus (JOSV) replicates without causing overt disease [18]. In contrast, BRBV, which is phylogenetically closest to OZV, fails to replicate in immunocompetent mice, and lethal infection models have been established only in IFN signaling-deficient mice [10]. Based on these observations, we established a lethal OZV infection model using B6 Ifnar1 KO mice. Intraperitoneal infection with 10^3^–10^4^ PFU of OZV-P17 reproducibly resulted in 100% mortality, providing a standardized lethal mouse model (Fig 1A). These findings suggest that OZV, similar to BRBV, is highly sensitive to innate immune responses and requires impaired IFN signaling to establish productive infection and disease. Interestingly, a small number of mice survived infection at higher inoculation doses (10^5^–10^6^ PFU) (Fig 1A and 1B). In immunodeficient mouse models, viral lethality does not always correlate with inoculation dose [19]. In Japanese encephalitis virus infection, high-dose inoculation has been suggested to induce stronger innate immune responses, thereby attenuating disease severity [25]. In addition, IFN-α/β and IFN-γ pathways have been shown to act synergistically to control BRBV infection [26]. In the present study, OZV-infected B6 Ifnar1 KO mice exhibited robust induction of IFN-γ (Fig 2G), suggesting that high-dose infection may enhance immune pathways independent of type I IFNs, thereby contributing to survival in a subset of animals. Further studies are required to clarify the immune mechanisms underlying this phenomenon.

Several thogotoviruses, including THOV, DHOV, BRBV, and OZV, have been reported to cause human disease, although the number of documented cases remains limited [1,9,10,12]. Febrile illness appears to be a common clinical feature among these infections [1,9,10,12]. In the present study, among organs examined in OZV-infected B6 Ifnar1 KO mice, the liver and spleen exhibited pronounced viral replication with comparatively lower viral burdens in the lung, kidney, and serum (Fig 2A and 2B). Marked elevation of inflammatory cytokine expression was observed particularly in the liver (Fig 2G), and hepatocellular necrosis was a prominent pathological feature accompanied by increased ALT and AST, which serve as liver injury biomarkers (Fig 2C and 2I). Dysregulated inflammatory responses in the absence of type I IFN signaling may contribute to excessive liver pathology, consistent with reports indicating that type I IFN plays a critical role in the negative regulation of inflammation [27]. In line with these observations in mice, highly elevated liver injury biomarkers were noted in the human OZV case, as well as in reported BRBV infection cases [9,10,12]. Although viral loads in the liver were reportedly low in postmortem samples from the fatal human case of OZV infection after 26 days of hospitalization, liver injury markers peaked at day 2 of hospitalization [12], suggesting that viral replication-associated liver injury may contribute to clinical manifestations during the acute to middle phase of infection in humans. Notably, focal hepatocellular necrosis, as observed in our OZV and previous BRBV mouse models, has been commonly found in THOV- and DHOV-infected mice [10,26,28–30], suggesting that hepatic involvement may represent a shared clinical feature within the genus *Thogotovirus*. Taken together, this mouse model demonstrates prominent hepatic infection with injuries during lethal OZV infection and may provide a pathological context for the liver enzyme elevation reported in the human case. Of note, the only reported fatal human case of OZV infection was associated with severe myocarditis, with viral RNA detected in myocardial cells [12]. In the present mouse model, no viral replication or pathological lesions were detected in the hearts (Fig 2A and 2B; S8 Fig), indicating that cardiac involvement is not a prominent feature under our experimental conditions. However, given the scarcity of human cases, it remains unclear whether myocarditis represents a typical manifestation of OZV infection in humans.

Currently, there are no approved antiviral drugs or vaccines for OZV. Using this established lethal mouse model, we demonstrated that oral administration of T-705 completely protected B6 Ifnar1 KO mice from lethal OZV infection and effectively suppressed viral replication and pathological changes (Fig 3B–H). Furthermore, immunization with an inactivated whole-virus vaccine induced neutralizing antibodies and conferred complete protection against lethal challenges (Fig 4C and 4D). This study demonstrates the utility of this mouse model for potential antiviral treatment and vaccine evaluation against OZV infection.

In conclusion, we have established a lethal OZV infection model in B6 Ifnar1 KO mice that partially recapitulates hepatic injury induced by OZV infection and enables evaluation of antiviral drugs and vaccines. Under our experimental conditions, myocarditis, which was observed in the human fatal case, was not detected, indicating that further investigation is required to clarify the involvement of OZV in this specific disorder. Nevertheless, our model will facilitate further studies on OZV pathogenesis and contribute to the development of effective countermeasures against this emerging tick-borne virus.

## Materials and methods

### Ethical statement

All animal experiments were reviewed and approved by the Institutional Animal Care and Use Committee of Hokkaido University (approval number: 24-0034) and were conducted in accordance with the committee’s guidelines.

### Animals

BALB/c mice were obtained from Japan SLC, Inc. (Shizuoka, Japan). B6-Ifnar1 knockout (KO) mice (C57BL/6J-Ifnar1<em1>) were generated using CRISPR/Cas9 genome editing, following the strategy outlined in S3 Fig, and were supplied from a colony maintained at the Central Institute for Experimental Medicine and Life Science. All animal experiments were performed at the International Institute for Zoonosis Control, Hokkaido University, under the approval described above. Humane endpoints were defined as a 25% loss of the maximum body weight or an inability to access food or water due to the severity of the disease.

### Cells and viruses

Vero E6 cells were maintained in Dulbecco’s Modified Eagle Medium (DMEM, FUJIFILM Wako Pure Chemical Corporation, Osaka, Japan) supplemented with 10% fetal bovine serum (FBS) at 37℃ with 5% CO_2_. The OZV Ibaraki/O10-S/2022 strain [12] was passaged 17 times in Vero E6 cells to generate OZV-P17. Virus stocks were stored at −80°C until use.

### Virus titration by plaque assay

Monolayers of Vero E6 cells in 24-well plates were inoculated with 10-fold serial dilutions of samples in DMEM supplemented with 10% FBS. After 1 h of absorption at 37℃, the inoculum was removed, and the cells were washed with phosphate-buffered saline (PBS). Cells were overlaid with 0.5% methylcellulose in Eagle’s Minimum Essential Medium (EMEM, Nissui Pharmaceutical Co., Ltd., Tokyo, Japan) supplemented with 2% FBS, GlutaMAX (Gibco, Thermo Fisher Scientific, Waltham, MA, USA), and 1% Antibiotic-Antimycotic (Gibco). After incubation for 4 days at 37℃, cells were fixed with 10% formalin and stained with 1% crystal violet. Plaques were counted, and viral titers were defined as plaque-forming units (PFU) per milliliter.

### In vivo infection and pathogenicity analysis in B6 Ifnar1 KO mice

B6 Ifnar1 KO mice (seven-week-old) were inoculated with 10^2^, 10^3^, 10^4^, 10^5^, or 10^6^ PFU of OZV-P17 in 50 µl of PBS via i.p. or s.c. routes under anesthesia. Body weight changes and survival were monitored for 16 days. For virological and pathological analyses, brain, heart, lung, liver, spleen, kidney, and serum samples were collected from mice at 1, 3, and 5 days after the i.p. inoculation with 10^4^ PFU of OZV-P17. Tissues were homogenized in PBS for titration and RNA extraction or fixed in 10% buffered formalin for histopathological analysis.

### Reverse transcription-quantitative PCR (RT-qPCR)

Total RNA was extracted from tissue homogenate using TRIzol LS reagent (Invitrogen, Thermo Fisher Scientific, Waltham, MA, USA) and Direct-zol RNA Miniprep kit (Zymo Research, Irvine, CA, USA). Quantification of OZV viral RNA was performed using THUNDERBIRD Probe One-step qRT-PCR Kit (Toyobo, Osaka, Japan) with specific primers (forward: 5’-TCATCGACCACAACGCAGAA-3’; reverse: 5’-GGTCCCATCTTTGAGGGTGG-3’) and a probe (5’-FAM-GCGTCCATTGTGATGGCAGCC-BHQ-3’) targeting the NP segment. Reactions were performed under the following conditions: 10 min at 50℃, 1 min at 95℃, and 45 cycles of 15 sec at 95℃ and 45 sec at 60℃ using QuantStudio 7 Flex Real-Time PCR System (Thermo Fisher Scientific). Viral RNA copy numbers were calculated by the standard curve method. For analysis of inflammatory gene expression, One Step TB Green PrimeScript RT-PCR Kit II (Perfect Real Time) (Takara Bio, Shiga, Japan) and specific primer sets targeting IFN-γ, IL-1β, IL-6, TNF-α, and GAPDH were used (S1 Table [31–34]). Reactions were performed under the following conditions: 5 min at 42℃, 10 sec at 95℃, and 50 cycles of 5 sec at 95℃ and 20 sec at 60℃ using LightCycler 96 (Roche, Basel, Switzerland). Relative gene expression levels in infected mice relative to mock mice were calculated using the ΔΔCt method, with GAPDH used as the reference gene.

### Histopathological analysis

Collected mouse tissues were fixed in 10% buffered formalin for 7 days and then embedded in paraffin. Sections were stained with hematoxylin and eosin (HE) and for immunohistochemical analysis. For antigen retrieval, the sections were treated with citrate buffer for 3 min using a pressure cooker. Then, the sections were treated with 0.3% H_2_O_2_ in methanol for 15 min at room temperature to inactivate endogenous peroxidase. Histofine mouse staining kit (Nichirei Biosciences Inc., Tokyo, Japan) was used for blocking and secondary antibody reactions according to the manufacturer’s protocol. Serum from a surviving OZV-infected BALB/c (Figure S2) mouse (1:1,000) was used as a primary antibody for OZV detection. Images were acquired using NanoZoomer S60v2 (Hamamatsu Photonics, Shizuoka, Japan) and analyzed with NDP. view2 software (Hamamatsu Photonics).

### Quantification of the histopathological lesions in the liver

QuPath v0.6.0 [35,36], an open software for bioimage analysis, was used to quantify histopathological lesions in the liver. Whole liver tissue sections were segmented by threshold-based methods. To exclude edge-related nonspecific signals, image analysis was restricted to regions located more than 200 µm from the tissue boundaries. To ensure consistent analysis across slides with various staining intensities, normal regions, vascular regions, artifact regions, focal necrosis, and parenchymal degeneration were annotated in all mock and infected samples (Fig 2). Based on these annotations, a pixel classifier was trained using a machine-learning-based tool implemented in QuPath. After classification, vascular and artifact regions were excluded from further analysis. The relative areas of focal necrosis and parenchymal degeneration were then calculated as histopathological regions.

### Serum biochemical analysis

Serum biochemical parameters were analyzed using Multi Roter I Preventive Care Panel (Zoetis Japan, Tokyo, Japan) on VetScan VS2 (Zoetis Japan). Briefly, serum (100 µl) was loaded onto the panel according to the manufacturer’s instructions. Serum samples showing hemolysis-related errors were excluded from the analysis.

### Antiviral treatment with T-705

B6 Ifnar1 KO mice (seven-week-old) were inoculated with 10^4^ PFU of OZV-P17 in 50 µl of PBS via i.p. route under anesthesia. T-705 treatment was initiated immediately after infection. Mice were orally administered 200 µl of 0.5% methylcellulose with or without T-705 (Angene International, Nanjing, China) at doses of 100 or 300 mg/kg/day, twice daily until 8 dpi. Body weight and survival were monitored for 20 days. For virological and histopathological analyses, brain, heart, lung, liver, spleen, kidney, and serum samples were collected at 5 dpi from mice treated with vehicle or T-705 (300 mg/kg/day), and subjected to virus titration, RT-qPCR, and histopathological analysis as described above.

### Inactivated vaccine preparation and immunization

Culture supernatants of Vero E6 cells infected with the OZV-P17 strain were clarified by centrifugation at 1,800 × *g* for 15 min. Formalin was added to a final concentration of 0.1%, and the mixture was incubated for 7 days at 4℃ to inactivate the virus. The inactivated virus was purified by ultracentrifugation through a 20% sucrose cushion at 28,000 rpm for 2 h at 4℃, and the pellet was resuspended in PBS. Complete inactivation was confirmed by the absence of cytopathic effect after three serial passages in Vero E6 cells. Total protein concentration of the purified inactivated virions was measured using Pierce BCA Protein Assay Kits (Thermo Fisher Scientific). Vaccines were prepared by mixing inactivated virions containing 1.5 μg or 7.5 μg of total protein with an equal volume of AddaS03 (InvivoGen, San Diego, CA, USA) [37,38]. B6 Ifnar1 KO mice (ten-week-old) were immunized subcutaneously twice at 24 days interval. Serum samples were collected 24 days and 16 days after each immunization, and binding antibody titer and neutralizing antibody titers were measured.

### ELISA

ELISA plates were coated overnight at 4°C with lysates of OZV-P17-infected Vero E6 cells diluted in coating buffer (Medicago Inc., Quebec City, QC, Canada). Plates were blocked with 5% skim milk in PBS containing 0.01% Tween 20 (PBST) for 1 h at room temperature. Serially diluted serum (1:100 to 1:1,638,400) in 5% skim milk in PBST was added and incubated for 1 h at 37°C. After washing three times with PBST, HRP-conjugated anti-mouse IgG (1:3,000, Nacalai Tesque Inc., Kyoto, Japan) in 5% skim milk in PBST was added and incubated for 1 h at 37°C. After 3-times washing with PBST, 1-Step Ultra TMB-ELISA Substrate Solution (Thermo Fisher Scientific) was added, and the reaction was stopped by adding 2M H_2_SO_4_. Absorbance at 450 nm was measured using Infinite 200 PRO (Tecan Group Ltd., Männedorf, Switzerland). The endpoint titer was determined as the reciprocal number of the highest serum dilution at which the absorbance value exceeded the mean plus three standard deviations of blank samples.

### Neutralization assays

For BALB/c mice, serum was diluted in a 2-fold serial dilution from 1:5 to 1:160 in DMEM supplemented with 10% FBS and mixed with 50 PFU of OZV-P17 at an equal volume. For vaccinated B6 Ifnar1 KO mice, serum was diluted in a 2-fold serial dilution from 1:25 to 1:800 and mixed with 450 PFU of OZV-P17. After incubation for 1 h at 37°C, the mixtures were inoculated onto Vero E6 cell monolayers in 24-well plates. After 1 h absorption, the inocula was removed, and cells were overlaid with 0.5% methylcellulose in EMEM supplemented with 2% FBS, GlutaMAX, and 1% Antibiotic-Antimycotic. After incubation for 4 days at 37℃, cells were fixed with 10% formalin and stained with 1% crystal violet. The 50% plaque reduction neutralization test (PRNT_50_) titers were defined as the reciprocal number of the highest serum dilution resulting in greater than 50% reduction in plaque numbers compared with virus-only controls.

### Statistical analysis

All statistical analyses were performed using GraphPad Prism 10.4.2. Comparison between the two groups was conducted using the Mann-Whitney U test. Comparisons among multiple groups were performed using two-way analysis of variance (ANOVA) followed by Tukey’s multiple-comparison test. Statistical significance was defined as *, *P* < 0.05; **, *P* < 0.01; ***, *P* < 0.001; ****, *P* < 0.0001. Samples below the detection limit were statistically assigned a value equal to half of the detection limit for statistical analysis.

## Acknowledgments

We thank Dr. Yasuko Orba, Dr. Michihito Sasaki, Dr. Masashi Shingai, and Dr. Takuma Ariizumi for providing access to the equipment and for helpful advice on serum biochemical analysis. This study was supported in part by the Japan Society for the Promotion of Science (JSPS) KAKENHI under grant numbers 25K18810 and 25K09431; and the Japan Agency for Medical Research and Development (AMED) under grant numbers JP223fa627005 and JP24wm0225044. Schematic figures were created with BioRender.com.

## Author Contributions

**Conceptualization:** Rio Harada, Koshiro Tabata, Saito Shinji, Gabriel Gonzalez, Hirofumi Sawa, Yukari Itakura

**Data Curation:** Rio Harada, Yukari Itakura

**Formal Analysis:** Rio Harada

**Funding Acquisition:** Hirofumi Sawa, Yukari Itakura

**Investigation:** Rio Harada, Cassia Sousa Moraes, Atsuko Inoue, Chilekwa Frances Kabamba, Koshiro Tabata, Joshua Walter Kranrod, Aiko Ohnuma, Yukari Itakura

**Methodology:** Rio Harada, Yukari Itakura

**Project Administration:** Hirofumi Sawa, Yukari Itakura

**Resources:** Motohito Goto, Riichi Takahashi, Shuichi Osawa, Noriko Nagata, Mamoru Ito, Makoto Suematsu

**Supervision:** Hirofumi Sawa, Yukari Itakura

**Validation:** Rio Harada

**Visualization:** Rio Harada

**Writing – Original Draft Preparation:** Rio Harada

**Writing – Review & Editing:** Rio Harada, Cassia Sousa Moraes, Atsuko Inoue, Chilekwa Frances Kabamba, Koshiro Tabata, Motohito Goto, Riichi Takahashi, Joshua Walter Kranrod, Shuichi Osawa, Aiko Ohnuma, Saito Shinji, Gabriel Gonzalez, Noriko Nagata, Mamoru Ito, Makoto Suematsu, Hideki Hasegawa, William W. Hall, Hirofumi Sawa, Yukari Itakura

