## Supplemental for "A lethal mouse model of Oz virus infection reveals hepatic involvement and enables evaluation of antiviral and vaccine efficacy"

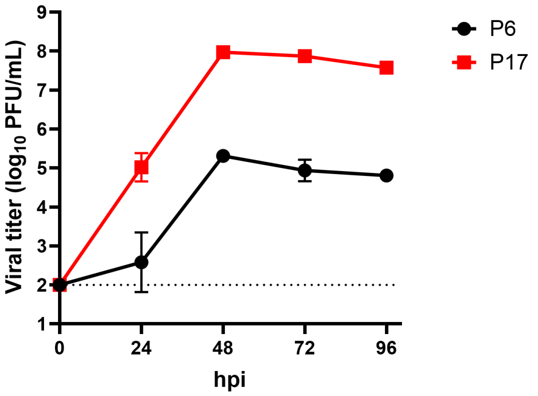


**S1 Fig. Growth curve of OZV before and after cell adaptation.** Vero E6 cells were infected with OZV-P6 and cell-adapted OZV-P17 at a multiplicity of infection of 0.01. Infectious viral titers were measured at the indicated time points. Data are presented as means ± SD of triplicate experiments. A dotted line indicates the lower limit of detection.

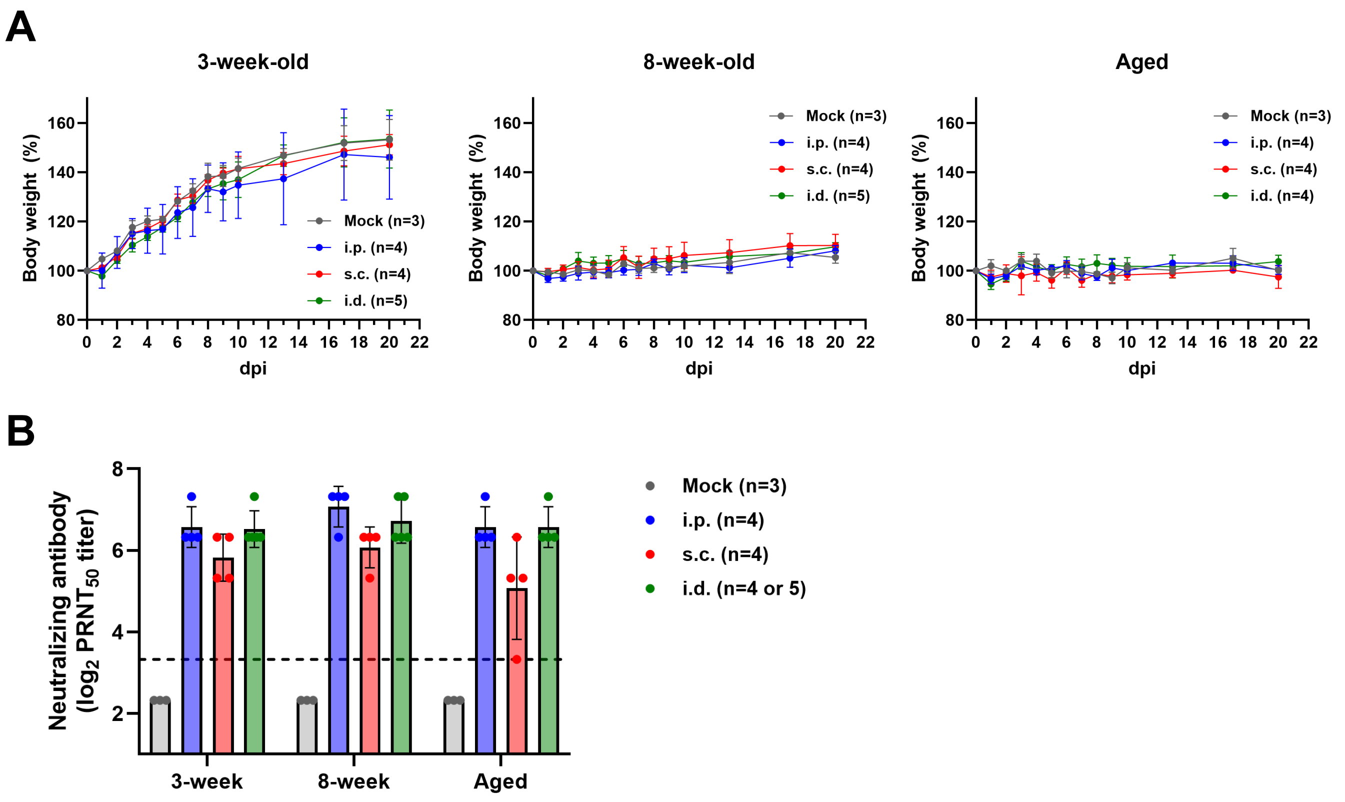


**S2 Fig. OZV-P17 infection in BALB/c mice.** Three-week-old, 8-week-old, and aged BALB/c mice were inoculated with $10^6$ PFU of OZV-P17 intraperitoneally (i.p.), subcutaneously (s.c.), or intradermally (i.d.) under anesthesia and monitored daily for 20 days. Body weight was measured at 0–10, 13, 17, and 20 days post infection (dpi). (A) Body weight changes were monitored until 20 dpi. (B) Serum samples were collected at 21 dpi, and neutralizing antibody titers were measured. A dotted line indicates the lower limit of detection. Data are presented as means ± SD in panels A and B, and each dot represents an individual mouse in panel B.

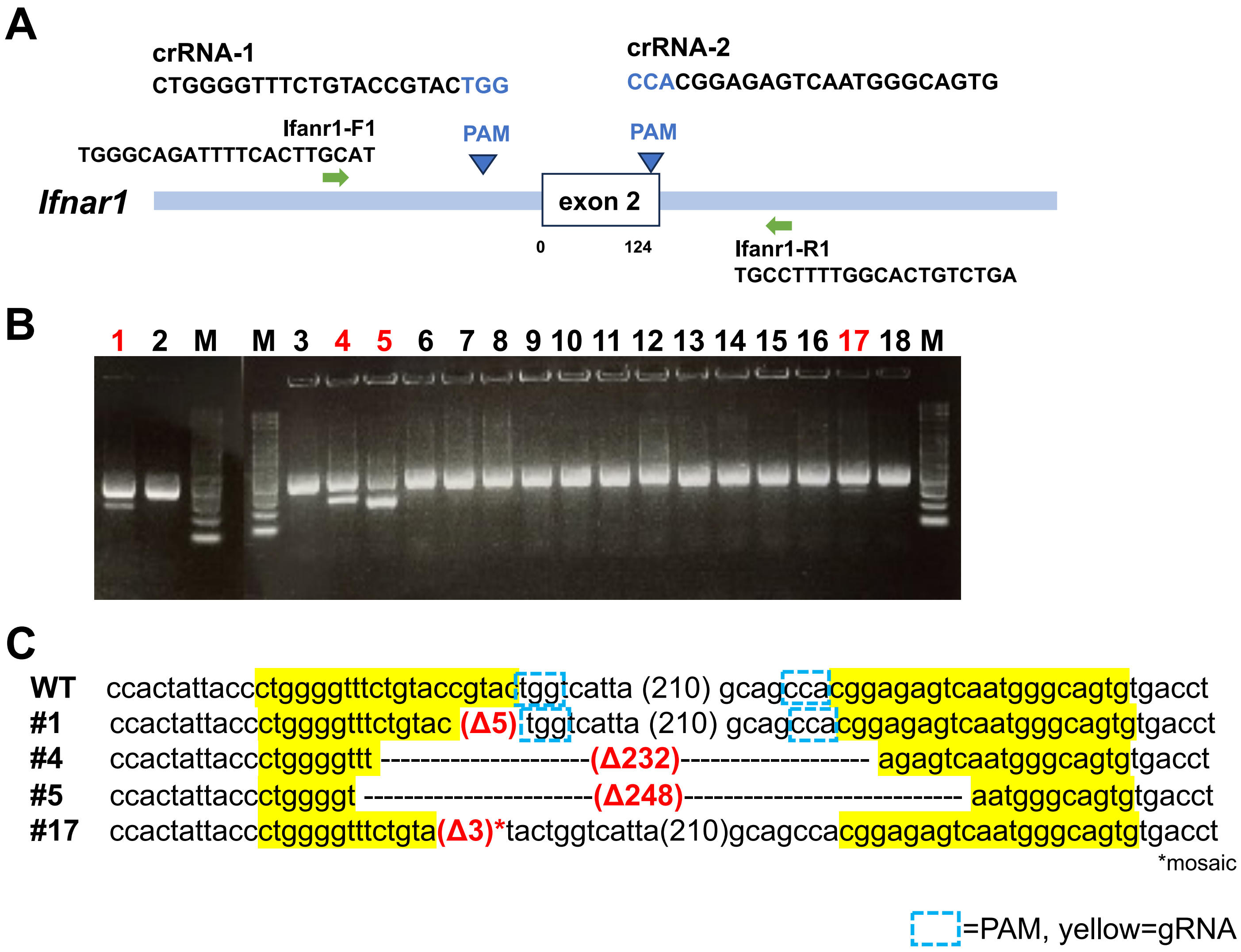


**S3 Fig. Generation of B6-Ifnar1 KO Mice Using CRISPR-Cas9.** (A) Schematic Diagram of the Ifnar1 Deficiency. (B) Agarose gel electrophoresis of PCR products amplified using primers Ifnar1-F1 and Ifnar1-R1, as shown in (A). (C) Sequence analysis of mutations around the protospacer adjacent motif (PAM; highlighted in blue) and the target sequence (highlighted in yellow), validated by Sanger sequencing. Electroporation [39] was performed to introduce the CRISPR–Cas9 construct into 95 fertilized C57BL/6J eggs. Following embryo transfer, 22 pups were obtained, of which 18 survived to weaning. PCR and sequence analyses identified four founder mice carrying large deletions. Among these, founder mouse #4, harboring a 232-bp deletion, was crossed with C57BL/6J mice to establish the B6-Ifnar1 KO strain (C57BL/6J-Ifnar1<em1>/Jic), which was used in all subsequent experiments.

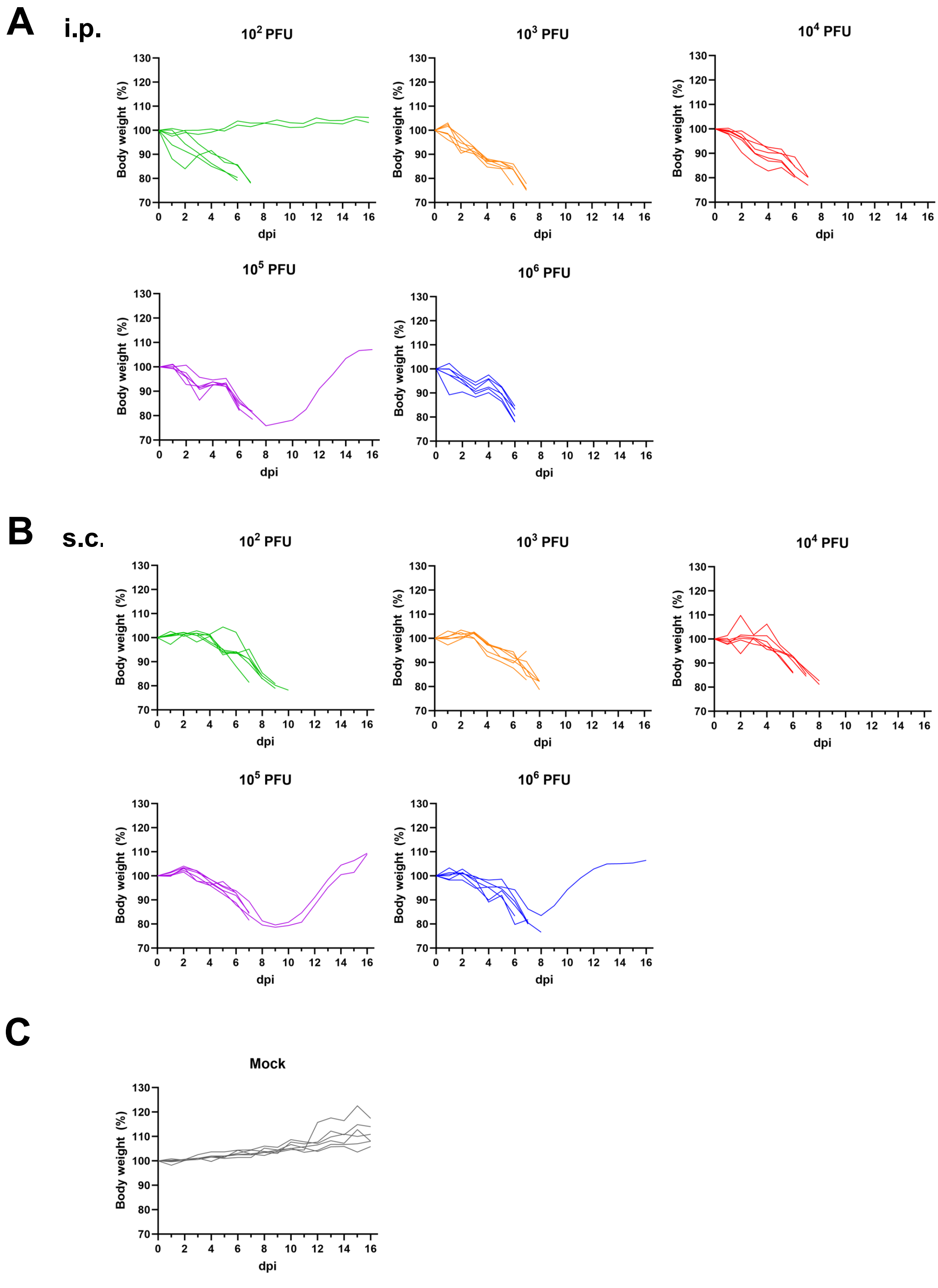


**S4 Fig. Body weight changes of an individual B6 Ifnar1 KO mouse infected with OZV-P17 (related to Fig 1).** B6 Ifnar1 KO mice (n=6) were inoculated with $10^2$, $10^3$, $10^4$, $10^5$, or $10^6$ PFU of OZV-P17 (A) intraperitoneally (i.p) or (B) subcutaneously (s.c.). (C) Mice in the mock group (n=6) received no treatment. Each line graph represents the body weight changes of an individual mouse.

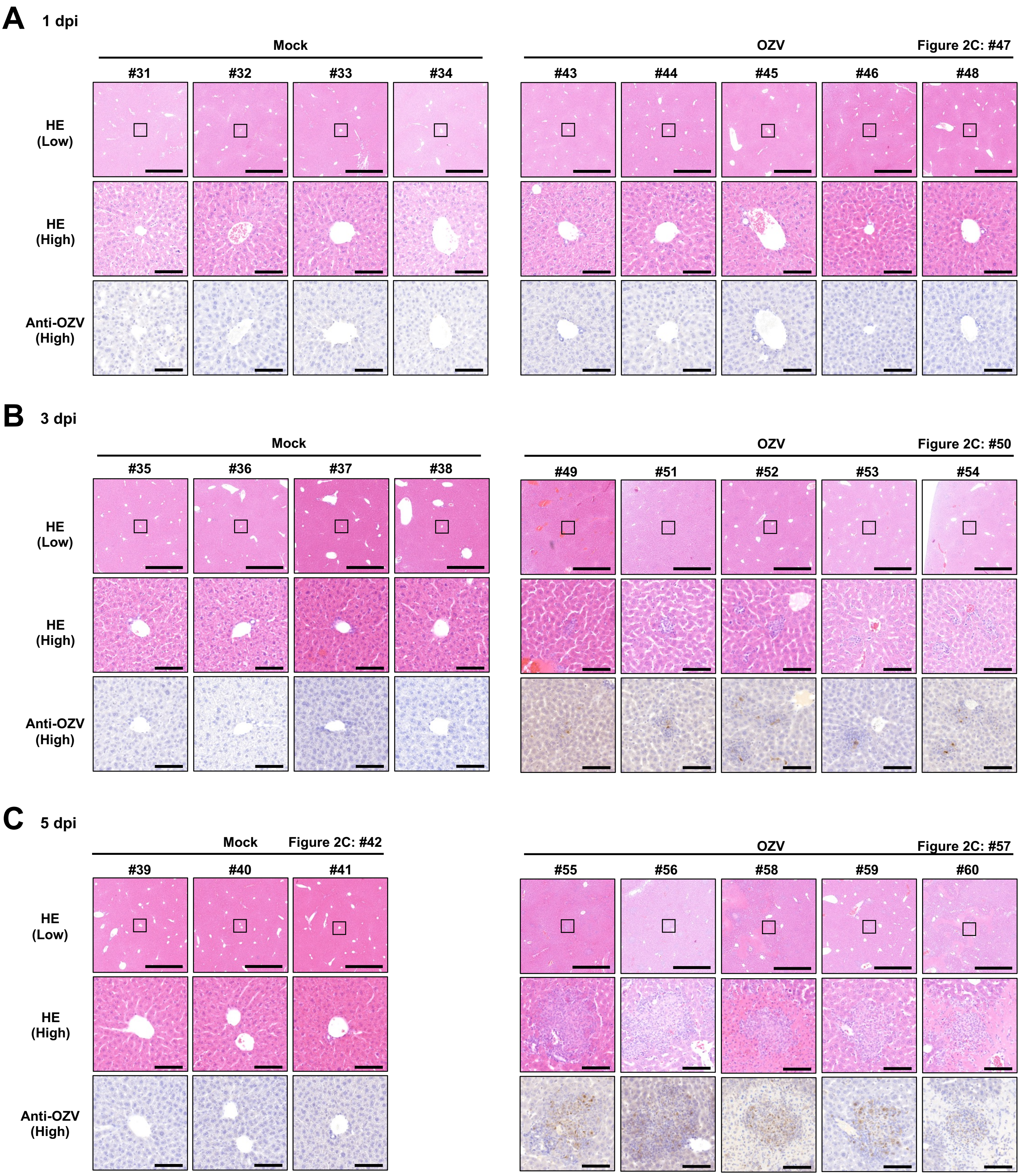


**S5 Fig. Histopathological analysis of the liver of B6 Ifnar1 KO mice infected with OZV-P17 (related to Fig 2C).** B6 Ifnar1 KO mice were inoculated intraperitoneally with PBS or $10^4$ PFU of OZV-P17. Images of hematoxylin and eosin (HE) staining and immunohistochemical detection of OZV in the liver at (A) 1, (B) 3, and (C) 5 dpi are shown. Scale bars represent 1 mm and 100 µm in the low- and high-magnification images, respectively. The boxed area in the low-magnification image is shown at high-magnification.

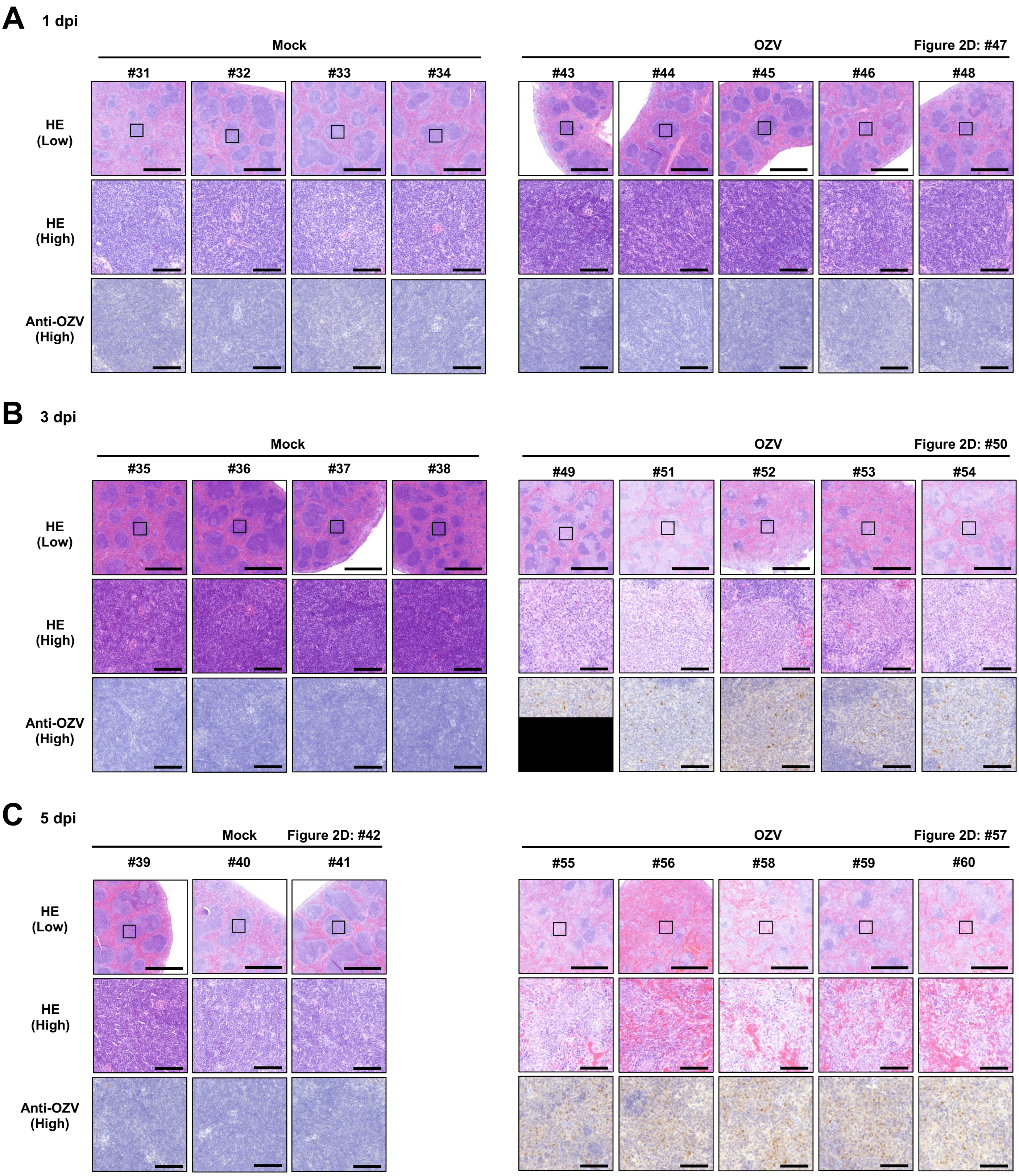


**S6 Fig. Histopathological analysis of the spleen of B6 Ifnar1 KO mice infected with OZV-P17 (related to Fig 2D).** B6 Ifnar1 KO mice were inoculated intraperitoneally with $10^4$ PFU of OZV-P17. Images of HE staining and immunohistochemical detection of OZV in the spleen at (A) 1, (B) 3, and (C) 5 dpi are shown. Scale bars represent 1 mm and 100 μm in the low- and high-magnification images, respectively. The boxed area in the low-magnification image is shown at high-magnification.

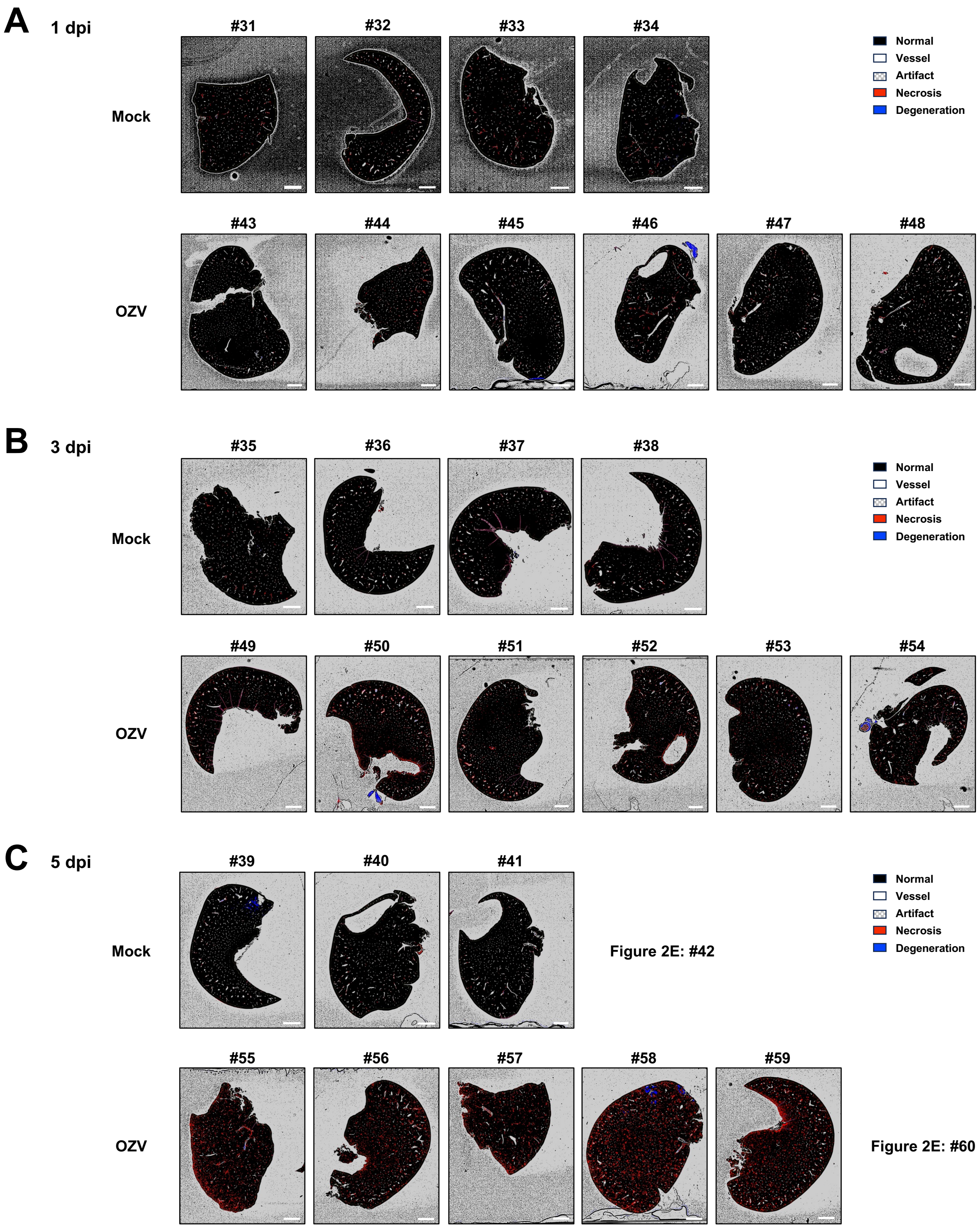


**S7 Fig. Lesion quantitative analysis of the liver of B6 Ifnar1 KO mice infected with OZV-P17 (related to Fig 2E and 2F).** B6 Ifnar1 KO mice were inoculated intraperitoneally with $10^4$ PFU of OZV-P17. Images of the quantitative analysis of the liver lesion at (A) 1, (B) 3, and (C) 5 dpi are shown. Scale bars represent 2 mm in the images. The boxed area in the low-magnification image is shown at high-magnification.

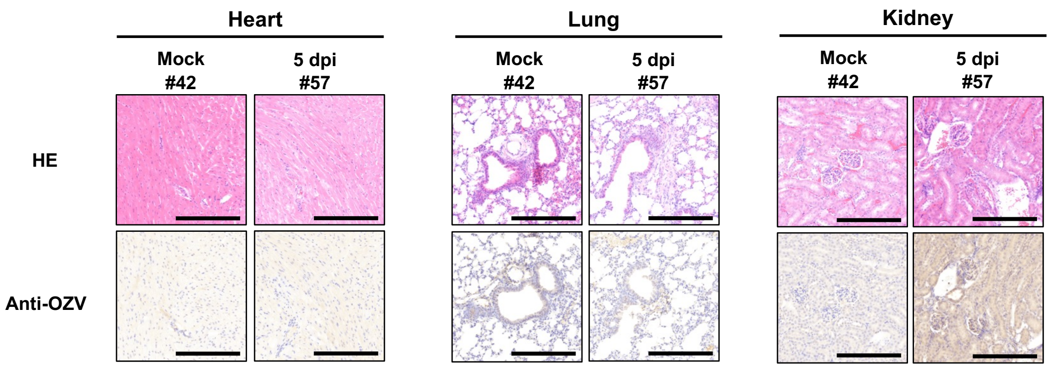


**S8 Fig. Histopathological analysis of the heart, lung, and kidney of B6 Ifnar1 KO mice infected with OZV-P17.** B6 Ifnar1 KO mice were inoculated intraperitoneally with $10^4$ PFU of OZV-P17. Representative images of HE staining and immunohistochemical detection of OZV in the heart, lung, and kidney at 5 dpi are shown. Scale bars represent 250 µm in the images.

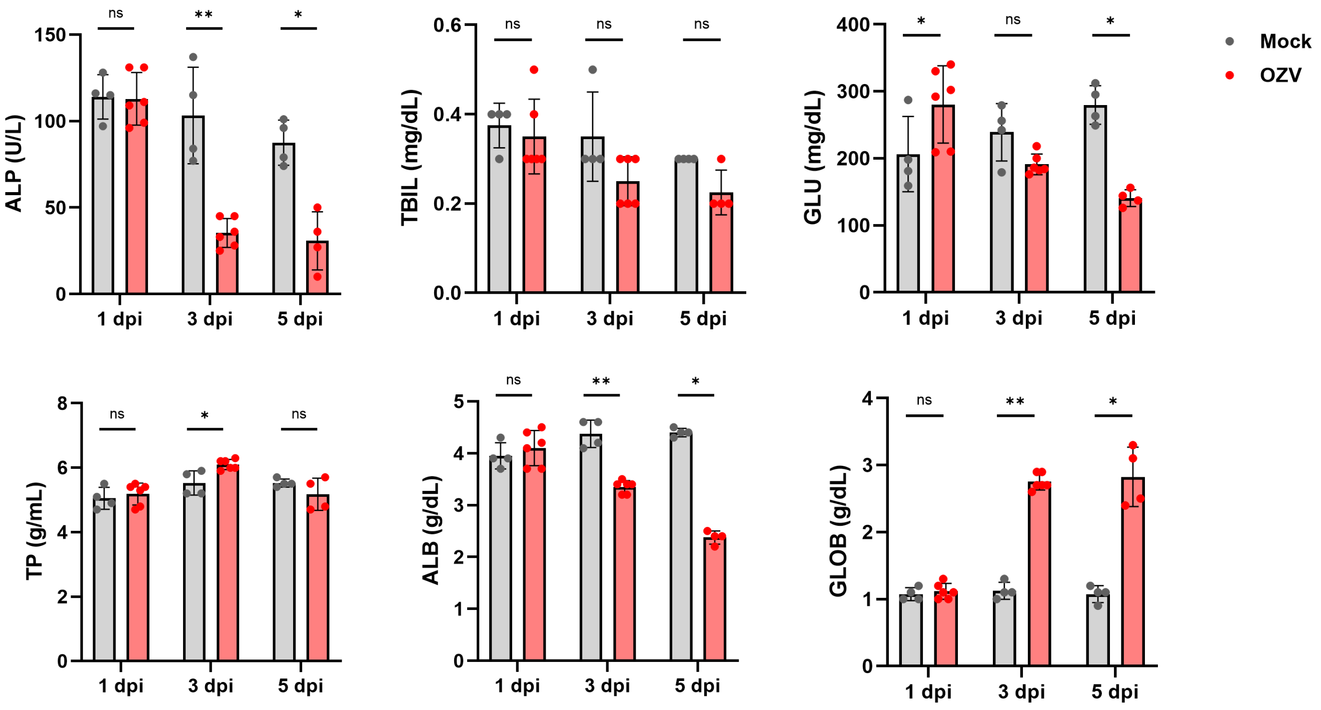


**S9 Fig. Serum biochemical analysis of B6 Ifnar1 KO mice infected with OZV-P17 (related with Fig 2I).** B6 Ifnar1 KO mice were inoculated intraperitoneally $10^4$ PFU of OZV-P17. Serum levels of alkaline phosphatase (ALP), total bilirubin (TBIL), glucose (GLU), total protein (TP), albumin (ALB), and globulin (GLOB) at each time point are shown. Data are presented as means ± SD, and each dot represents an individual mouse. Statistical analysis was performed using the Mann-Whitney U test: *, $P < 0.05$; **, $P < 0.01$.

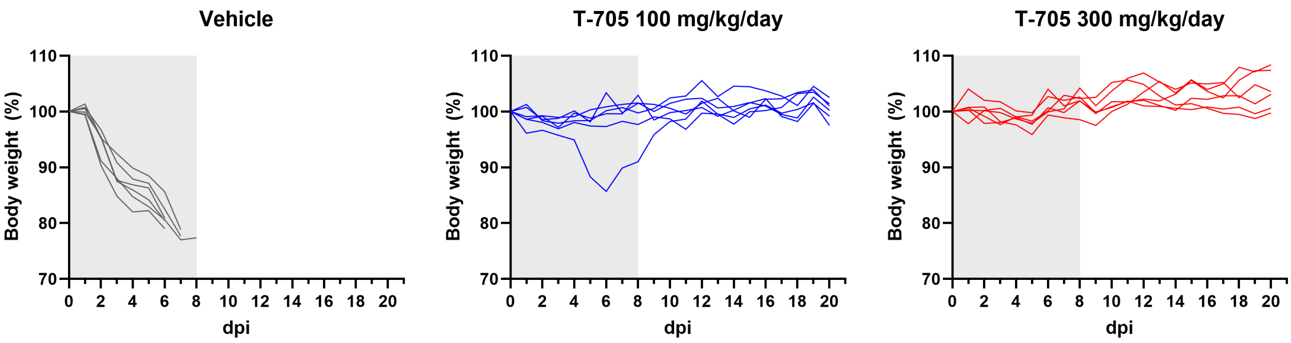


**S10 Fig. Body weight changes of an individual B6 Ifnar1 KO mouse treated with T-705 (related to Fig 3B).** B6 Ifnar1 KO mice (n=6) were intraperitoneally inoculated with 104 PFU of OZV-P17, and immediately administered orally with 100 or 300 mg/kg/day of T-705, or vehicle for 8 dpi. The treatment period is indicated by the gray shaded area. Each line represents body weight changes of an individual mouse.

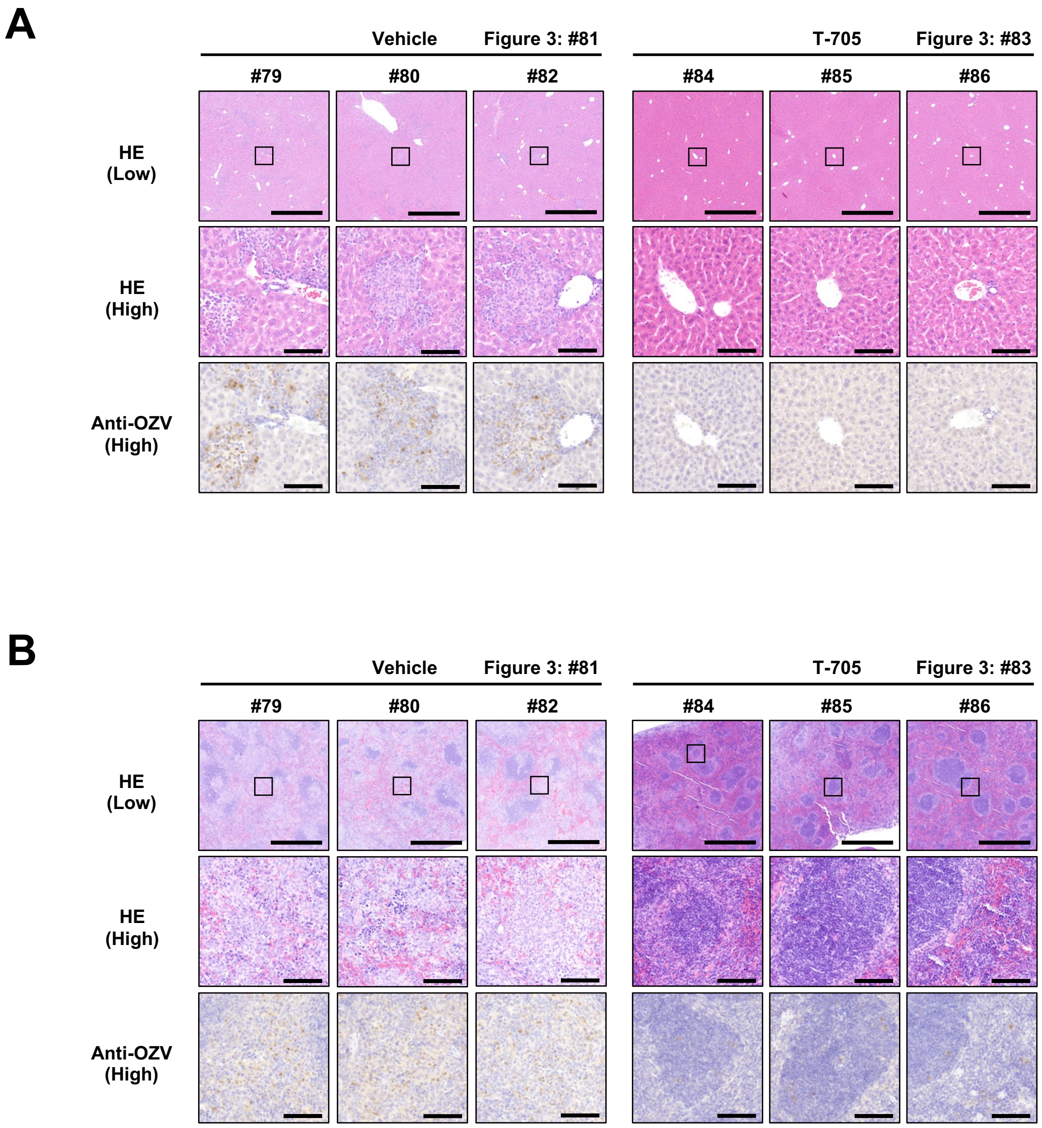


**S11 Fig. Histopathological analysis of the liver and spleen of B6 Ifnar1 KO mice treated with T-705 (related to Fig 3E and 3F).** B6 Ifnar1 KO mice were inoculated intraperitoneally with $10^4$ PFU of OZV-P17, and immediately administered orally with 300 mg/kg/day of T-705 or vehicle. Livers and spleens were collected at 5 dpi. Images of HE staining and immunohistochemical detection of OZV in the (A) liver and (B) spleen from an individual mouse are shown. Scale bars represent 1 mm and 100 µm in the low- and high-magnification images, respectively. The boxed area in the low-magnification image is shown at high-magnification.

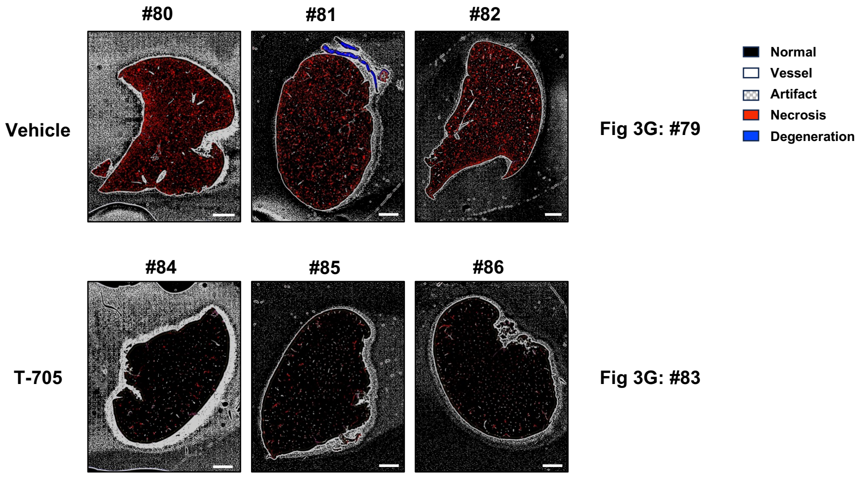


**S12 Fig. Lesion quantitative analysis of the liver of B6 Ifnar1 KO mice treated with T-705 (related to Fig 3G and 3I).** B6 Ifnar1 KO mice were inoculated intraperitoneally with $10^4$ PFU of OZV-P17, and immediately administered orally with 300 mg/kg/day of T-705 or vehicle. Livers were collected at 5 dpi. Images of the quantitative analysis of the liver lesion are shown. Scale bars represent 2 mm in the images. The boxed area in the low-magnification image is shown at high-magnification.

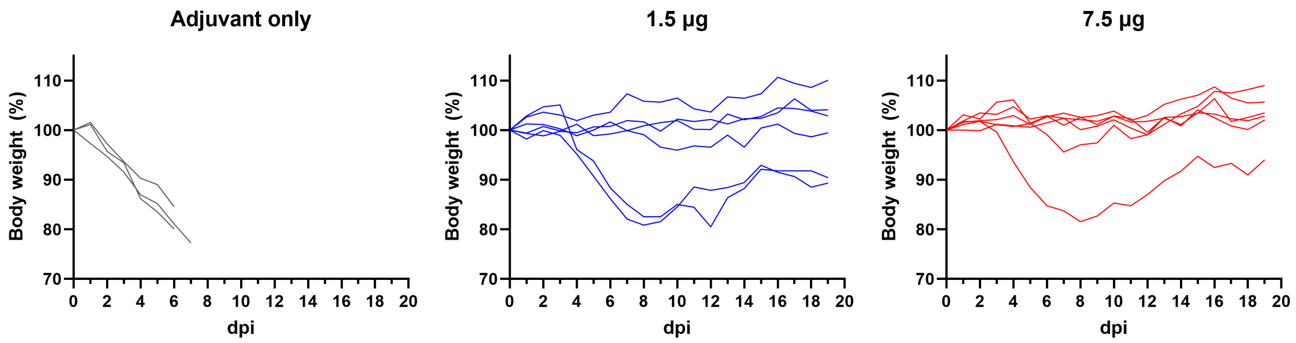


**S13 Fig. Body weight changes of an individual B6 Ifnar1 KO mouse in the vaccine challenge experiment (related to Fig 4D).** B6 Ifnar1 KO mice were immunized with 1.5 µg (n=6) or 7.5 µg (n=6) of antigen, or PBS (n=3), with adjuvant. Mice were challenged intraperitoneally with $10^4$ PFU of OZV-P17 three weeks after the boost immunization and monitored for 20 days. Each line represents body weight changes of an individual mouse.

**S1 Table. Mouse primers used for reverse transcription-quantitative PCR**

| Gene | Sequence of primer | | Source |
|---|---|---|---|
| INF-γ | Forward | 5'-TCAAGTGGCATAGATGTGGAAGAA-3' | Mohammed et al., 2011 |
| | Reverse | 5'-TGGCTCTGCAGGATTTTCATG-3' | |
| IL-1β | Forward | 5'-GAAATGCCACCTTTTGACAGTG-3' | Sun et al., 2016 |
| | Reverse | 5'-TGGATGCTCTCATCAGGACAG-3' | |
| IL-6 | Forward | 5'-TAGTCCTTCCTACCCCAATTTCC-3' | Zhao et al., 2022 |
| | Reverse | 5'-TTGGTCCTTAGCCACTCCTTC-3' | |
| TNF-α | Forward | 5'-CTGAACTTCGGGGTGATCGG-3' | Humbert-Claude et al., 2016 |
| | Reverse | 5'-GGCTTGTCACTCGAATTTTGAGA-3' | |
| GAPDH | Forward | 5'-AGGTCGGTGTGAACGGATTTG-3' | PrimerBank ID: 6679937a1 |
| | Reverse | 5'-TGTAGACCATGTAGTTGAGGTCA-3' | |
